# New method to reconstruct phylogenetic and transmission trees with sequence data from infectious disease outbreaks

**DOI:** 10.1101/069195

**Authors:** Don Klinkenberg, Jantien Backer, Xavier Didelot, Caroline Colijn, Jacco Wallinga

## Abstract

Whole-genome sequencing (WGS) of pathogens from host samples becomes more and more routine during infectious disease outbreaks. These data provide information on possible transmission events which can be used for further epidemiologic analyses, such as identification of risk factors for infectivity and transmission. However, the relationship between transmission events and WGS data is obscured by uncertainty arising from four largely unobserved processes: transmission, case observation, within-host pathogen dynamics and mutation. To properly resolve transmission events, these processes need to be taken into account. Recent years have seen much progress in theory and method development, but applications are tailored to specific datasets with matching model assumptions and code, or otherwise make simplifying assumptions that break up the dependency between the four processes. To obtain a method with wider applicability, we have developed a novel approach to reconstruct transmission trees with WGS data. Our approach combines elementary models for transmission, case observation, within-host pathogen dynamics, and mutation. We use Bayesian inference with MCMC for which we have designed novel proposal steps to efficiently traverse the posterior distribution, taking account of all unobserved processes at once. This allows for efficient sampling of transmission trees from the posterior distribution, and robust estimation of consensus transmission trees. We implemented the proposed method in a new R package *phybreak*. The method performs well in tests of both new and published simulated data. We apply the model to to five datasets on densely sampled infectious disease outbreaks, covering a wide range of epidemiological settings. Using only sampling times and sequences as data, our analyses confirmed the original results or improved on them: the more realistic infection times place more confidence in the inferred transmission trees.

**Author Summary:** It is becoming easier and cheaper to obtain whole genome sequences of pathogen samples during outbreaks of infectious diseases. If all hosts during an outbreak are sampled, and these samples are sequenced, the small differences between the sequences (single nucleotide polymorphisms, SNPs) give information on the transmission tree, i.e. who infected whom, and when. However, correctly inferring this tree is not straightforward, because SNPs arise from unobserved processes including infection events, as well as pathogen growth and mutation within the hosts. Several methods have been developed in recent years, but none so generic and easily accessible that it can easily be applied to new settings and datasets. We have developed a new model and method to infer transmission trees without putting prior limiting constraints on the order of unobserved events. The method is easily accessible in an R package implementation. We show that the method performs well on new and previously published simulated data. We illustrate applicability to a wide range of infectious diseases and settings by analysing five published datasets on densely sampled infectious disease outbreaks, confirming or improving the original results.

## Introduction

As sequencing technology becomes easier and cheaper, detailed outbreak investigation increasingly involves whole-genome sequencing (WGS) of pathogens from host samples [1]. These sequences can be used for studies ranging from virulence or resistance related to particular genes [1, 2], to the interaction of epidemiological, immunological and evolutionary processes on the scale of populations [3, 4]. If most or all hosts in an outbreak are sampled, it is also possible to use differences in nucleotides, i.e. single-nucleotide polymorphisms (SNPs), to resolve transmission clusters, individual transmission events, or complete transmission trees. With that information it becomes possible to identify high risk contacts and superspreaders, as well as characteristics of hosts or contacts that are associated with infectiousness and transmission [5, 6]. Much progress has been made in recent years in theory and model development, but existing methods typically include assumptions to address specific datasets, with fit-for-purpose code for data analysis. An easily accessible method with the flexibility to cover a wide range of infections is currently lacking, and would bring analysis of outbreak sequence data within reach of a much broader community.

The interest in easily applicable methods for sequence data analysis in outbreak settings is demonstrated by the community’s widespread use of the Outbreaker package in R [7-10]. However, the model in Outbreaker assumes that mutations occur at the time of transmission, which does not take the pathogen’s in-host population dynamics into account, nor the fact that mutations occur within hosts. The publications by Didelot et al [11] and Ypma et al [12] revealed that within-host evolution is crucial to relate sequence data to transmission trees, as is illustrated in Fig 1A: there are four unobserved processes, i.e. the time between subsequent infections, the time between infection and sampling, the pathogen dynamics within hosts, and mutation. The difference in sequences between host b and infector a result from all of these processes. As a result, a host’s sample can have different SNPs from his infector’s (Fig 1B: hosts a and b); a host can even be sampled earlier than his infector with fewer SNPs (Fig 1B: hosts a and c).

**Fig 1.**
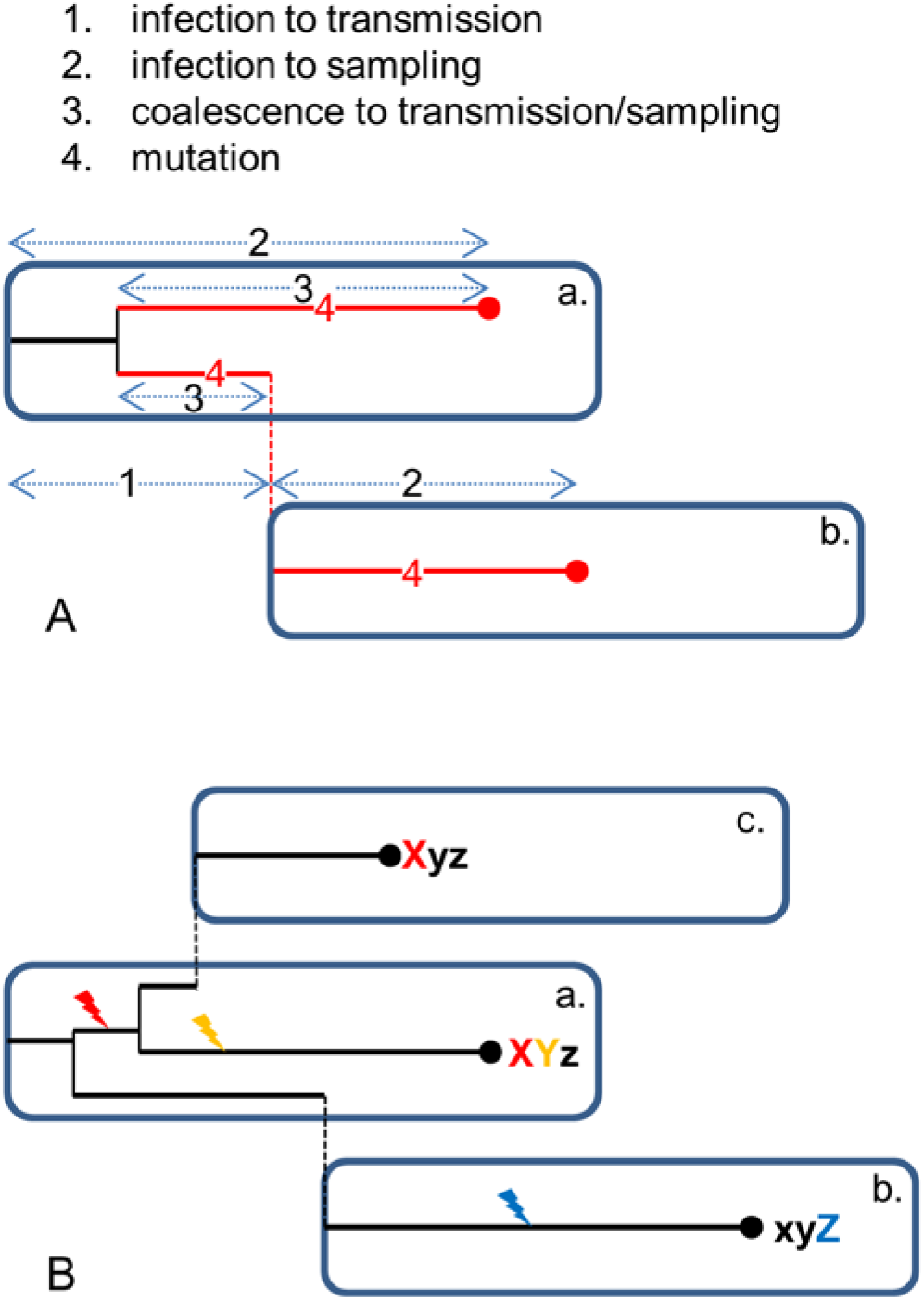
Sketch of stochastic processes involved in data generation process. (A) The four processes indicated by host a infecting host b. (B) Examples of resulting differences in sequences for host a infecting both hosts b and c.

Several recently published methods do allow mutations to occur within the host, but make other assumptions not fully reflecting the above-described process, such as using a phenomenological model for pairwise genetic distances [13], presence of a single dominant strain in which mutations can accumulate [14], or absence of a clearly defined infection time [15]. To take the complete process into account, Didelot et al [11] and Numminen et al [16] took a two-step approach: first, phylogenetic trees were built, and second, these trees were used to infer transmission trees. Didelot et al [11] used the software BEAST [17, 18] to make a timed phylogenetic tree, and used a Bayesian MCMC method to colour the branches such that changes in colour represent transmission events. Numminen et al [16] took the most parsimonious tree topology, and accounted for unobserved hosts by a sampling model (which is an additional complication). This two-step approach is likely to work better if the phylogenetic tree is properly resolved (unique sequences with many SNPs), but less so if there is uncertainty in the phylogenetic tree. However, also in that case construction of the phylogenetic tree is done without taking into account that only lineages in the same host can coalesce, and that these go through transmission bottlenecks during the whole outbreak. That is likely to result in incorrect branch lengths and consequently incorrect infection times.

Hall and Rambaut [19] implemented a method in BEAST for simultaneous inference of transmission and phylogenetic trees. BEAST allows for much flexibility when it comes to phylogeny and population dynamics reconstruction (for which it was originally developed [17, 18]), e.g. by allowing variation in mutation rates between sites in the genome, between lineages, and in time. However, datasets of fully observed outbreaks often do not contain sufficient information for reliable inference: they typically cover only a few months up to at most several years (as in Didelot et al [11], with tuberculosis) and do not contain many SNPs (usually of the same order of magnitude as the number of samples). A more important limitation is that the transmission model implemented in BEAST is rather specific: it allows for transmission only during an infectious period constrained by positive and negative samples, during which infectiousness is assumed to be constant. This may put prior constraints on the topology and order of events in the transmission and phylogenetic trees, which is undesirable if the primary aim is to reconstruct the transmission tree with little or no prior information about when hosts were infectious.

Previously, Ypma et al [12] had also developed a method for simultaneous inference of transmission and phylogenetic trees, albeit with rather specific assumptions on the within-host pathogen dynamics and the time and order of transmission events, and with no available implementation. However, their view on the phylogenetic and transmission trees was quite different. Instead of a phylogenetic tree with transmission events, they regarded it as a hierarchical tree. The top level is the transmission tree, with hosts having infected other hosts according to an epidemiological transmission model. The lower level consists of phylogenetic “mini-trees” within each host. A mini-tree describes the within-host micro-evolution. It is rooted at the infection time and has tips at transmission and sampling events. The complete phylogenetic tree then consists of all these mini-trees, connected through the transmission tree. That description allowed them to develop new MCMC updating steps, some changing the transmission tree, some the phylogenetic mini-trees.

We built further on that principle to reconstruct the transmission trees of outbreaks, in a new model and estimation method. The method requires data on pathogen sequences and sampling times. The model includes all four underlying stochastic processes (Fig 1A), each described in a flexible and generic way, such that we avoid putting unnecessary prior constraints on the order of unobserved events (Fig 1B). This allows for application of the method to a wide range of infectious diseases, including new emerging infections where we have little quantitative information on the infection cycle. The method is implemented in R, in a package called *phybreak*. We illustrate the performance of the method by applying it to both new and previously published simulated datasets. We demonstrate the range of applicability by applying the model to to five datasets on densely sampled infectious disease outbreaks, covering a wide range of epidemiological settings.

## Results

### Outline of the method

The method infers infection times and infectors of all cases in an outbreak. The data consist of sampling times and sequences of all cases, where some of the sequences may be empty if no sequence is available. Using simple models for transmission, sampling, within-host dynamics and mutation, samples are taken from the posterior distributions of model parameters and transmission and phylogenetic trees, by a Markov-Chain Monte Carlo (MCMC) method. The main novelty of our method lies in the proposal steps for the phylogenetic and transmission trees that are used to generate the MCMC chain. It makes use of the hierarchical tree perspective, in which the phylogenetic tree is described as a collection of phylogenetic mini-trees (one for each host), connected through the transmission tree (see Methods for details).

The posterior samples are summarized by medians and credible intervals for parameters and infection times, and by consensus transmission trees. Consensus transmission trees are based on the posterior support for infectors of each host, defined as the proportion of posterior trees in which a particular infector infects a host. The Edmonds’ consensus tree takes for each host the infector with highest support, and uses Edmonds’ algorithm to resolve cycle and multiple index cases [20], whereas the Maximum Parent Credibility (MPC) tree is the one tree among the posterior trees with maximum product of supports [19].

The models and parameters used for inference are as follows:

- transmission: assuming that all cases are sampled and the outbreak is over, the mean number of secondary infections must be 1. The transmission model therefore consists only of a Gamma distribution for the generation interval, i.e. the time interval between a primary and a secondary case. The model contains two parameters: the shape *a_G_*, which we fixed at 3 during our analyses, and the mean *m_G_*, which is estimated and has a prior distribution with mean *μ_G_* and standard deviation *σ_G_*. In an uninformative analysis, *μ_G_* = 1, and *σ_G_* = ∞.
- sampling: the sampling model consists of a Gamma distribution for the sampling interval, which is the interval between infection and sampling of a case. The model contains two parameters: the shape *a_S_*, which is fixed during the analysis, and the mean *m_S_*, which is estimated and has a prior distribution with mean *μ_S_* and standard deviation *σ_S_*. In an uninformative analysis, *μ_S_* = 1, and *σ_S_* = ∞; in a naïve analysis we additionally set *a_S_* = 3.
- within-host dynamics: The within-host model describes a linearly increasing pathogen population size *w*(*τ*) = *rτ*, at time *τ* since infection of a host. The slope *r* has a Gamma distributed prior distribution with shape *a_r_* and rate *b_r_*. In an uninformative analysis, *a_r_* = *b_r_* = 1.
- The mutation model is a site-homogeneous Jukes-Cantor model, with per-site mutation rate *μ*. The prior distribution for log(*μ*) is uniform.

### Analysis of the newly simulated datasets

We generated new simulated datasets were generated with the above model, in a population of 86 individuals and a basic reproduction number *R*_0_ = 1.5, to obtain 25 datasets of 50 cases. Parameters were *a_G_* = *a_S_* = 10, = *m_S_* = *r* = 1, *μ* = 10^-4^ and sequence length 10^4^, resulting in 1 genome-wide mutation per mean generation interval of one year.

Table 1 shows some summary measures on performance of the method (see S1 Results for additional measures and results for more simulations). Sampling a single chain of 25,000 MCMC cycles took about 30 minutes on a 2.6 GHz CPU (Linux). Four sets of results are shown: one with all parameters fixed at their correct value, and three with different levels of prior knowledge on *m_S_* only: informative with correct mean, uninformative, and informative with incorrect mean. The top of the table shows effective sample sizes (ESSs) for *μ* and *m_S_* and for the infection times to evaluare mixing of continuous parameter samples. To evaluate mixing across and within chains of infectors per host, we tested for differences between the chains and for dependency within the chains by Fisher’s exact tests: the proportion of accepted tests (*P* > 0.05) is a measure of mixing. The MCMC mixing is generally good for tree inference and model parameters, as most ESSs are above 200 and an expected 95% of Fisher’s tests is accepted; the only exceptions are the within-host parameter *r* (ESSs between 100 and 200, S1 Results), and *m_S_* with an uninformative prior.

**Table 1.** Performance on 25 newly simulated datasets of 50 cases, with shape parameters *a_S_* = *a_G_* = 10.

| | | Level of prior information on $m_S$ | | |
| --- | --- | --- | --- | --- |
|  | Reference <sup>a</sup> | Informative<br>Correct <sup>b</sup> | Uninformative <sup>c</sup> | Informative<br>Wrong <sup>d</sup> |
| **MCMC sampling** |  |  |  |  |
| Continuous parameter samples (95% interval of ESS) |  |  |  |  |
| $\mu$ | 6520 ; 9763 | 631 ; 1853 | 226 ; 1327 | 548 ; 1764 |
| $m_S$ | | 270 ; 319 | 31 ; 111 | 309 ; 789 |
| $t_{inf}$ | 1499 ; 8378 | 765 ; 4437 | 212 ; 982 | 425 ; 4314 |
| Infectors (% Fisher's exact tests accepted) |  |  |  |  |
| *between chains* | 98.6% | 98.3% | 98.7% | 98.6% |
| *autocorrelation* | 96.1% | 97.0% | 94.9% | 96.9% |
| **Tree inference** |  |  |  |  |
| Infection times (coverage: % of 95% CIs containing the true value) |  |  |  |  |
|  | 95.8% | 96.2% | 96.8% | 44.6% |
| Infectors (number correct/number identified) |  |  |  |  |
| *Edmonds'* | 34.9/50 | 34.6/50 | 34.3/50 | 34.5/50 |
| *MPC* | 33.4/50 | 32.4/50 | 32.6/50 | 30.7/50 |
| *>50% support* | 27.9/33.0 | 28.3/34.0 | 28.1/34.1 | 21.6/24.4 |
| *>80% support* | 15.2/15.7 | 15.2/15.8 | 15.4/16.0 | 8.2/8.4 |

Results are based on two MCMC chains of 25,000 samples each; ESS, effective sample size; CI, credible interval; MPC, maximum parent credibility. <sup>a</sup> $m_G, m_S, r = 1$ ; <sup>b</sup> $\mu_S = 1, \sigma_S = 0.1$ ; <sup>c</sup> $\mu_S = 1, \sigma_S = \infty$ ; <sup>d</sup> $\mu_S = 2, \sigma_S = 0.1$

The bottom part of Table 1 shows the results on tree inference. Infection times (using all MCMC samples) are well recovered if the mean sampling interval does not have a strong incorrect prior. For this simulation scenario, consensus transmission trees contained almost 70% (35 out of 50) correct infectors, as determined by counting infectors and resolving multiple index cases and cycles in the tree (Edmonds’ method [20]) and slightly fewer when choosing the maximum parent credibility (MPC) tree [19] among the 50,000 posterior trees. Infectors with high support are more likely correct: 82% (28 out of 34) are correct if the support is above 50%, and 96% (15.4 out of 16) are correct if the support is above 80%. These numbers are similar in smaller outbreaks, and lower if sampling and generation interval distributions are wider (S1 Results). Using prior information on the mean sampling interval did not improve on this, but if prior information is incorrect, fewer hosts have a strongly supported infector, which makes inference more uncertain. In conclusion, the method is fast and efficient if applied to simulated data fitting the model. In that case, no informative priors are needed for transmission tree inference.

### Analysis of previously published simulated data

We applied the method to previously published outbreak simulations [19]. Briefly, a spatial susceptible-exposed-infectious-recovered (SEIR) model was simulated in a population of 50 farms, with a latent period (exposed) of two days and a random infectious period with mean 10 days and standard deviation 1 day, at the end of which the farm was sampled. Two mutation rates were used, either *Slow Clock* or *Fast Clock*, equivalent to 1 or 50 genome-wide mutations per generation interval of one week, respectively.

Table 2 shows performance of the method with naïve and informative prior information on the sampling interval distribution (see S1 Results for uninformative). Effective sample sizes are a bit smaller than with the new simulations, but in most cases still good for infection times, whereas sampling of infectors was excellent. The low variance of the sampling interval distribution caused some problems in efficient sampling of *m_S_* because of its high correlation with the associated infection times. This is best seen in the ESS of *m_S_* and infection times in the uninformative *Slow Clock* analysis (S1 Results), but it also causes problems in the burn-in phase if inference starts with parameter values far from their actual values (not shown). Posterior median mutation rates are slightly higher than used for simulation, which could be due to different rates for transition and transversion in the simulation model [19].

**Table 2.** Performance on 25 published simulated datasets in populations of size 50 [19].

|  | *Slow Clock simulations* |  | *Fast Clock simulations* |  |
| --- | --- | --- | --- | --- |
| **Prior information** | **Naïve <sup>a</sup>** | **Informative <sup>b</sup>** | **Naïve <sup>a</sup>** | **Informative <sup>b</sup>** |
| **MCMC sampling** |  |  |  |  |
| Continuous parameter samples (95% interval of ESS) |  |  |  |  |
| $\mu$ | 225 ; 505 | 355 ; 979 | 24 ; 710 | 62 ; 702 |
| $m_S$ | 44 ; 114 | 37 ; 88 | 88 ; 932 | 79 ; 390 |
| $t_{inf}$ | 170 ; 1475 | 197 ; 608 | 170 ; 2064 | 236 ; 2571 |
| Infectors (% Fisher's exact tests accepted) |  |  |  |  |
| *between chains* | 95.9% | 97.3% | 87.3% | 96.8% |
| *autocorrelation* | 94.2% | 96.7% | 81.7% | 94.8% |
| **Parameter inference** (95% interval of posterior medians) |  |  |  |  |
| $\log_{10}(\mu)$ | -4.86 ; -4.66 | -4.93 ; -4.77 | -3.29 ; -3.17 | -3.20 ; -3.16 |
| $m_G$ | 2.1 ; 4.9 | 3.6 ; 5.7 | 4.2 ; 6.4 | 4.7 ; 6.1 |
| $m_S$ | 6.6 ; 10.3 | 11.3 ; 12.8 | 9.9 ; 15.5 | 11.6 ; 12.6 |
| $r$ | 0.40 ; 1.6 | 0.13 ; 0.59 | 0.49 ; 2.6 | 0.24 ; 1.3 |
| **Tree inference** |  |  |  |  |
| Infection times (coverage: % of 95% CIs containing the true value) |  |  |  |  |
|  | 76.6% | 97.6% | 95.4% | 94.7% |
| Infectors (number correct/number identified) |  |  |  |  |
| *Edmonds'* | 28.8/49.3 | 30.7/49.3 | 31.8/49.3 | 45.3/49.3 |
| *MPC* | 25.1/49.3 | 30.2/49.3 | 29.8/49.3 | 45.3/49.3 |
| *>50% support* | 12.9/14.3 | 25.4/31.4 | 22.6/28.7 | 45.0/48.7 |
| *>80% support* | 3.0/3.1 | 18.8/20.2 | 4.4/4.8 | 41.0/42.2 |

Results are based on two MCMC chains of 25,000 samples each. The mean outbreak size was 49.3 cases; ESS, effective sample size; CI, credible interval; MPC, maximum parent credibility. <sup>a</sup>
$a_S = 3, \mu_S = 1, \sigma_S = \infty$ ; <sup>b</sup> $a_S = 144, \mu_S = 12, \sigma_S = 1$

Consensus trees with uninformative and informative settings were almost as good as in the original publication [19], in which spatial data were used and in which it was known that there was a latent period and that hosts could not transmit after sampling. In the *Slow Clock* simulations about 62% of infectors were correct, and in the *Fast Clock* simulations about 92%. Infection times were also well recovered in most cases, but not in the uninformative *Slow Clock* analysis (S1 Results). In the naïve analyses, the *Slow Clock* consensus trees are only slightly less good (but mixing of the chain much better), whereas the *Fast Clock* consensus trees become much worse, with only 65% of infectors correct. In conclusion, the method performs well if applied to data simulated with a very different model. Good prior information on the variance of the sampling interval can significantly improve transmission tree inference, especially if the genetic data contain many SNPs.

### Analysis of published datasets

We finally applied the method to five published datasets on outbreaks of *Mycobacterium tuberculosis* (Mtb, [11]), Methicillin-resistant *Staphylococcus aureus* (MRSA, [21]), Foot-and-mouth disease (FMD2001 and FMD2007, [12, 22-24]), and H7N7 avian influenza (H7N7, [19, 25-27]).

The results for the four smaller datasets are shown in Table 3, which shows that mixing of the MCMC chains was generally good. Fig 2 shows the Edmond’s consensus trees (full details in S1 Results), with each host’s estimated infection time and most likely infector, colour coded to indicate posterior support. Fig 3 shows one sampled tree for each dataset (from the posterior set of 50,000), matching the MPC consensus tree topology.

**Fig 2.**
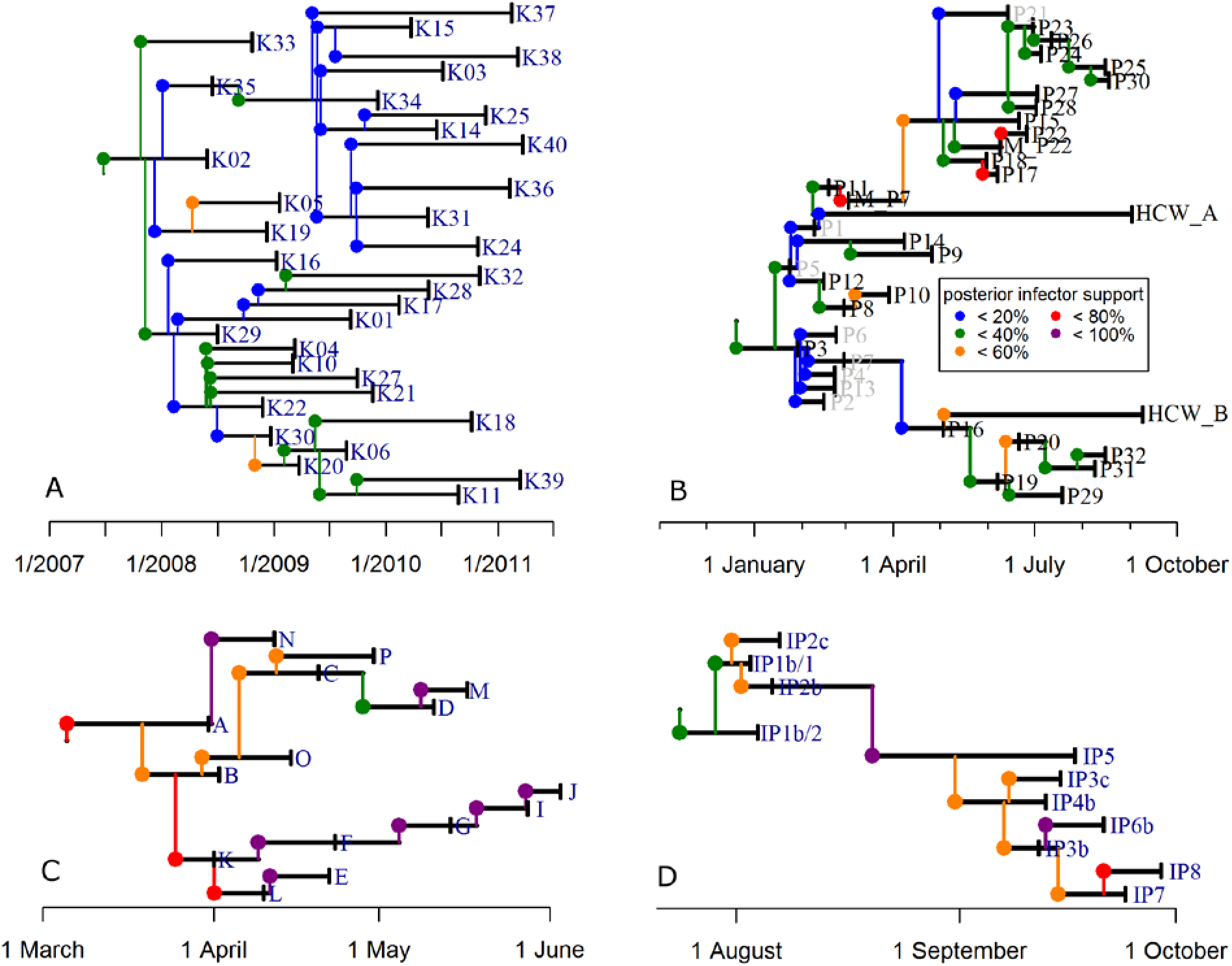
Consensus Edmonds’ transmission trees for four of the five analysed datasets. Vertical bars indicate sampling days, coloured links indicate most likely infectors, with colours indicating the posterior support for that infector. (A) Mtb data [11]; (B) MRSA data [21]; (C) FMD2001 data [22]; (D) FMD2007 data [23].

**Fig 3.**
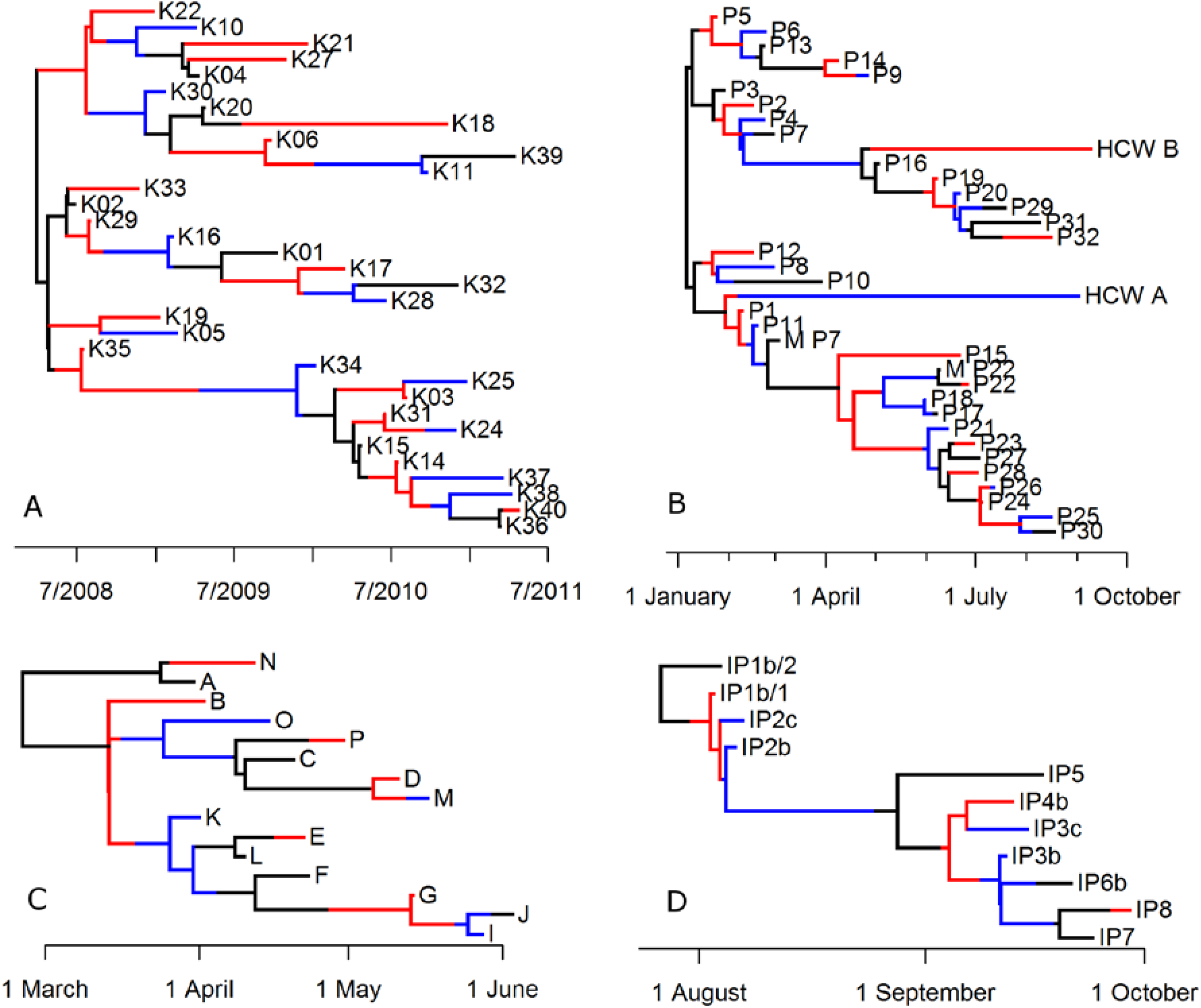
Consensus MPC transmission and phylogenetic trees for four of the five analysed datasets. Each tree is one posterior sample matching the MPC tree topology. Colours are used to indicate the hosts in the transmission tree: connected branches with identical colour are in the same host, and a change of colour along a branch is a transmission event. (A) Mtb data [11]; (B) MRSA data [21]; (C) FMD2001 data [22, 24]; (D) FMD2007 data [23, 24].

**Table 3.**
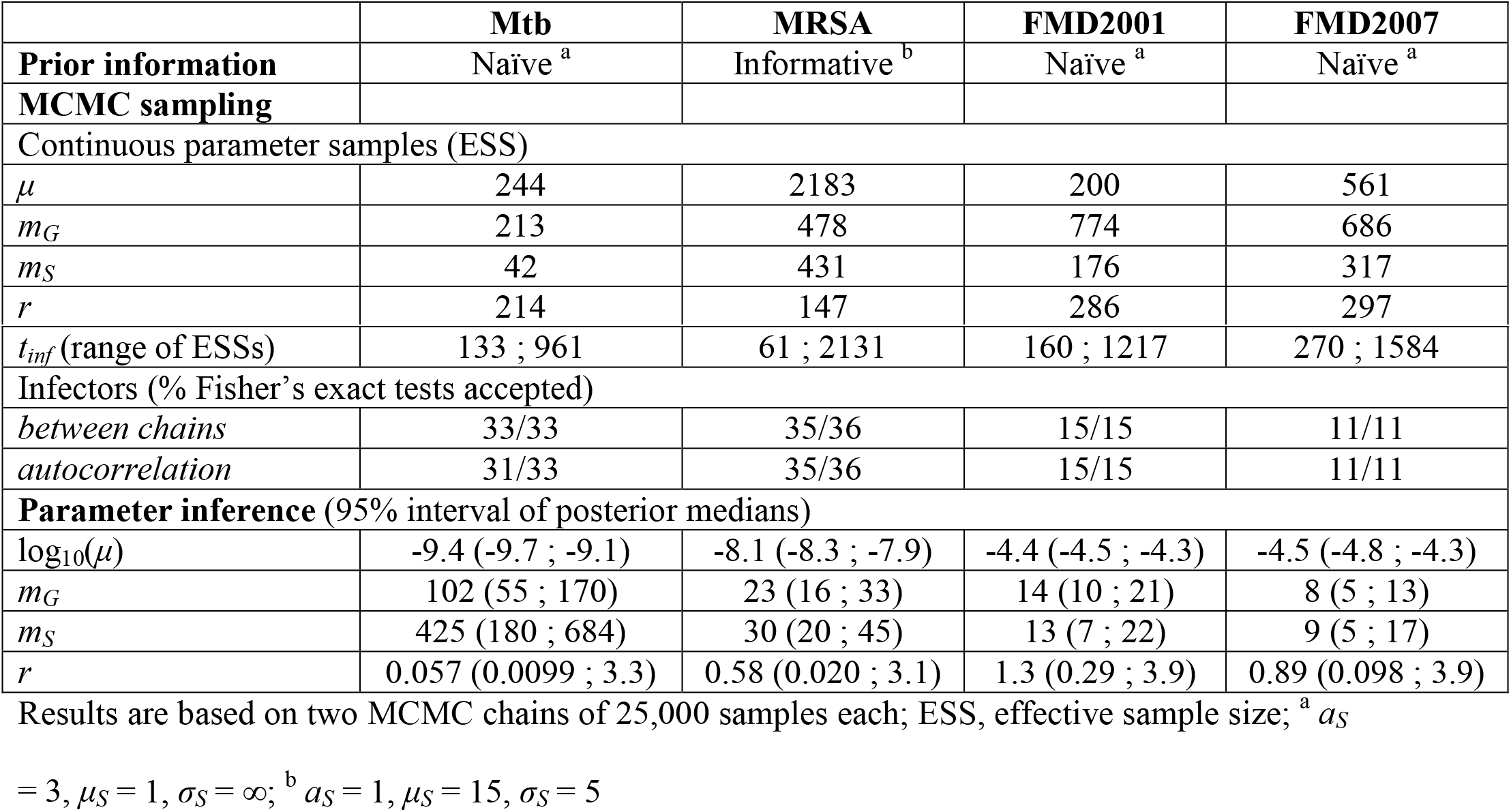
Summary statistics for four published datasets.

The Mtb data were analysed with naïve prior information, which resulted in a median sampling interval of 425 days (similar to estimated incubation times [28]), a median generation interval of 102 days, and a mutation rate equivalent to 0.3-1.3 mutations per genome per year, as estimated before [29, 30]. The Edmonds’ censensus transmission tree (Fig 2a) shows low support for most infectors, which is mainly a reflection of the low number of SNPs. However, the same index case K02 and three clusters as identified in Didelot et al [11] are distinguished: one starting with K22, one with K35, and the remaining cases starting with K16 or infected by the index case. The main difference compared to the original analysis lies in the shape of the phylogenetic tree and the estimated infection times (Fig 3a). Whereas the topology is very similar, the timing of the branching events is different: in the original tree, internal branches decrease in length when going from root to tips, consistent with a coalescent tree based on a single panmictic population. By taking into account the fact that coalescent events take place within individual hosts, our analysis shows branch lengths that are more balanced in length across the tree. Importantly, this results in a more recent dating of root of the tree: early 2008 (Fig 3a) instead of early 2004 [11].

The MRSA data were analysed with an informative prior for the mean sampling interval *m_S_* and a shape parameter *a_S_* based on data on the intervals between hospitalisation and the first positive sample. The estimated mutation rate is similar to literature estimates [31, 32], but the posterior median *m_S_* of 30 days is considerably higher than the prior mean of 15 days (Table 3). This may be explained by the two health-care workers (HCW_A and HCW_B), which have very long posterior sampling intervals that were not part of the data informing the prior (Edmonds’ consensus tree, Fig 2b). In contrast with the original analysis, we now identify a transmission tree rather than only a phylogenetic tree, resulting in the observation that the two health-care workers may not have infected any patient in spite of their long infection-to-sampling interval. Almost all transmission events with low support (<20%) involved unsequenced hosts. Three of them were identified as possible infector, in the initial stage of the outbreak, when only few samples were sequenced. This indicates that some unsequenced hosts may have played a role in transmission, but that it is not clear which. Finally, a major difference between our results and those in the original publication is the shape of the phylogenetic tree and the dating of the tree root: around 1st January (Fig 3b) instead of 1st September the year before [21].

Analysis of the FMD2001 and FMD2007 datasets resulted in posterior sampling intervals with means of 13 and 9 days, respectively, close to the 8.5 days estimated from epidemic data [33]. Generation intervals were about the same (Table 3). Both datasets contained more SNPs than the Mtb and MRSA data, with unique sequences for each host and higher mutation rates, similar to published rates in FMD outbreak clusters [34]. This resulted in equal Edmonds’ and MPC consensus transmission trees, and much higher support for most infectors (Figs 2cd, 3cd). Our transmission tree is almost identical to the one from Ypma et al [12], who used a closely related method but did not allow for transmission after sampling. When comparing to the analysis of these data by Morelli et al [24], the topologies of the phylogenetic trees (Fig 3cd) match the topologies of the genetic networks (Fig S18 in [24]), but the transmission trees are quite different. The main differences are that in the FMD2001 outbreak, they identify farm B as the infector of C, E, K, L, O, and P; and in the FMD2007 outbreak, they have IP4b infecting IP3b, IP3c, IP6b, IP7, and IP8. Differences are likely the result of their use of the spatial data [24]. Use of additional data is expected to improve inference, although their estimates of infection-to-sampling intervals (about 30 days) were unrealistically long.

The H7N7 dataset was analysed with the sequences of the three genes HA, NA, and PB2 separately, and combined; with informative priors for both the mean sampling and mean generation intervals. Five parallel chains were run, and mixing was generally good (Table 4); it took about 7 hours on a 2.6GHz CPU to obtain 25,000 unthinned samples in a single chain. Analysis of the three genes combined resulted in a posterior median *m_S_* of 8.5 days, slightly longer than the 7 days on which the informative prior was based [35], and longer than in the separate analyses. The mean generation time was shorter than the prior mean: 3.9 days with all genes. We also calculated the parsimony scores of the posterior sampled trees, defined as the minimum numbers of mutations on the trees that can explain the sequence data [36], and compared these with the numbers of SNPs in the data (Table 4). It appeared that with the genes separately analysed, parsimony scores were 6-13% higher than the numbers of SNPs, indicating some homoplasy in the phylogenetic trees (this was not seen with any of the other datasets). The analysis of all genes together resulted in parsimony scores of 18% higher than the number of SNPs. The estimated mutation rates are among the highest estimates for Avian Influenza Virus, as already noted before in earlier analyses of the same data [25, 37]. Fig 4 shows the Edmonds’ consensus tree in generations of infected premises, indicating locations and inferred infection days (full details in S1 Results). Without the use of location data, there is a large Limburg cluster, a Central cluster including two sampled Limburg cases, and a small Limburg cluster of three cases with an exceptionally long generation time (about 8 lines from the bottom). A closer look at the sequences makes clear that the first of these cases (L22/34) has 3 SNPs different from assigned infector G4/11, and 4 SNPs different from cases in the large Limburg cluster. Using geographic data as in earlier analyses [19, 27] will probably place these cases within that cluster.

**Fig 4.**
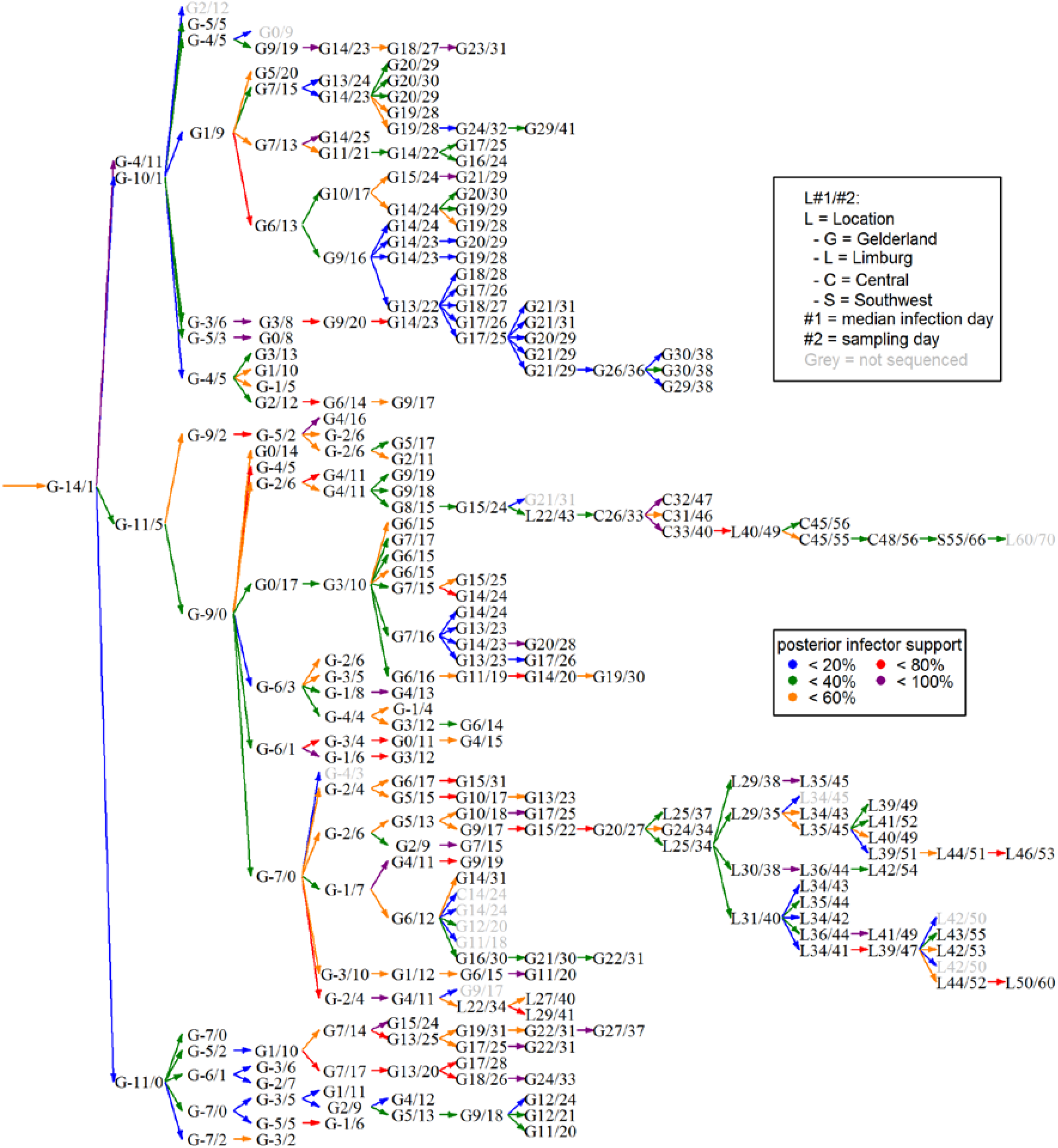
Consensus Edmonds’ transmission tree for the H7N7 dataset [19, 25, 27]. Infected premises are (not uniquely) coded by location (as in [19]), median posterior infection day, and sampling day. Coloured arrows indicate most likely infectors, with colours indicating the posterior support for that infector.

**Table 4.** Summary statistics for H7N7 dataset.

|  | Sequenced gene |  |  |  |
| --- | --- | --- | --- | --- |
|  | HA | NA | PB2 | All genes |
| **MCMC sampling** |  |  |  |  |
| Continuous samples <sup>a</sup> |  |  |  |  |
| $\mu$ | 1428 | 1366 | 2036 | 798 |
| $m_G$ | 854 | 838 | 1039 | 720 |
| $m_S$ | 330 | 240 | 311 | 232 |
| $r$ | 119 | 108 | 82 | 70 |
| $t_{inf}$ (range) | 400 ; 4873 | 409 ; 3920 | 538 ; 4225 | 82 ; 793 |
| Infectors <sup>b</sup> |  |  |  |  |
| *between chains* | 93.8% | 92.9% | 95.9% | 93.8% |
| *autocorrelation* | 97.1% | 97.1% | 95.9% | 97.9% |
| **Parameter inference <sup>c</sup>** |  |  |  |  |
| $\log_{10}(\mu)$ | -4.41 (-4.52 ; -4.31) | -4.42 (-4.54 ; -4.31) | -4.54 (-4.65 ; -4.44) | -4.50 (-4.56 ; -4.43) |
| $m_G$ | 3.6 (3.1 ; 4.2) | 3.4 (2.9 ; 4.0) | 3.8 (3.3 ; 4.4) | 3.9 (3.4 ; 4.5) |
| $m_S$ | 7.5 (6.7 ; 8.5) | 7.6 (6.6 ; 8.6) | 7.5 (6.7 ; 8.5) | 8.5 (7.6 ; 9.5) |
| $r$ | 0.21 (0.024 ; 1.5) | 0.25 (0.023 ; 1.5) | 0.085 (0.0026 ; 0.74) | 0.35 (0.029 ; 1.4) |
| **Phylogenetic inference** |  |  |  |  |
| Tree parsimony scores <sup>c</sup> | 102 (101 ; 103) | 83 (82 ; 85) | 100 (99 ; 101) | 313 (312 ; 315) |
| #SNPs in data | 90 | 73 | 94 | 257 |

Results are based on five MCMC chains of 25,000 samples each, with $a_S = 10$ ; $\mu_S = 7$ ; $\sigma_S = 0.5$ ;
$a_G = 3$ ; $\mu_G = 5$ ; $\sigma_G = 1$ ; $a_r = b_r = 1$ . SNP = single nucleotide polymorphism. <sup>a</sup> effective sample
sizes; <sup>b</sup> fraction of Fisher's exact tests with $P > 0.05$ ; <sup>c</sup> medians and 95% credible intervals

## Discussion

We developed a new method to reconstruct outbreaks of infectious diseases with pathogen sequence data from all cases in an outbreak. Our aim was to have an easily accessible and widely applicable method. For ease of access, we developed efficient MCMC updating steps which we implemented in a new R package, *phybreak*. We tested the method on newly simulated data, previously published simulated data, and published datasets. Our model is fast: 25,000 iterations took roughly 30 minutes with the Mtb and MRSA datasets of about 30 hosts, and 7 hours with the full three-genes H7N7 dataset in 241 hosts. Two chains with 50,000 posterior samples proved sufficient (measured by ESS) for tree inference (infectors and infection times) and most model parameters with simulated and published data. The package contains functions to enter the data, to run the MCMC chain, and to analyse the output, e.g. by making consensus trees and plotting these (as in Fig 3).

We tested the method on five published datasets, with outbreaks of viral and bacterial infections, and in diverse settings of open and closed populations, in human and veterinary context. The method performed well on all datasets in terms of MCMC chain mixing and tree reconstruction. With uninformative priors on mean sampling intervals and mutation rates, we obtained estimates that were all very accurate when compared to literature, and the inferred transmission trees seemed as good, or even better when considering estimated infection times. With two datasets (MRSA and H7N7) we included some prior information on sampling and/or generation intervals, which mainly affected the inferred infection times, but not so much the transmission trees.

For wide applicability, we kept the underlying model simple without putting prior constraints on the order of unobserved events such as infection and coalescence times. Four submodels with only one or two parameters each were used for sampling, transmission, within-host pathogen dynamics, and nucleotide substitution. The sampling model, a gamma distribution for the interval between infection and sampling, has a direct link to inferred infection times, and is the model for which it is most likely that prior information is available from epidemiological data in the same or other outbreaks. We used simulated data to study the effect of uninformative or incorrect prior information on shape parameter *a_S_* and mean *m_S_*. It appears that an incorrect *a_S_* or an incorrect informative prior for *m_S_* does reduce accuracy of inferred infection times. However, consensus trees are hardly affected, at least if the number of SNPs is in the order of the number of hosts as we saw in the actual datasets (Table 1 and Table 2 *Slow Clock*). Only the precision of consensus trees is reduced, i.e. there are fewer inferred infectors with high support. Results with the *Fast Clock* simulations did show a significant reduction in consensus tree accuracy. In that case, there are so many SNPs that the phylogenetic tree topology and times of coalescent nodes are almost fixed; then, too much variance in sampling intervals (low *a_S_*) results in many incorrect placements of infection events on that tree. Possibly, with so many SNPs it could be more efficient to first make the phylogenetic tree, and then use that tree to infer transmission events [11, 16], but it is questionable whether genome-wide mutation rates are ever so high that this becomes a real issue [38].

The submodel for transmission is relevant for transmission tree inference in describing the times between subsequent infection events. Transmission is modelled as a homogeneous branching process, implicitly assuming that there was a small outbreak in a large population, with a reproduction number (mean number of secondary cases per primary case) of 1. Our approach assumes that everyone in the outbreak was known, which is a potential limitation, as even with good surveillance, contact tracing, and case identification, there is always the possibility that some infectors are not known to outbreak investigators. If all, or almost all, infectors are in the data, the generation interval distribution reflects the course of infectiousness, separating the cases in time along the tree. Apart from not taking heterogeneity across hosts into account (an extension we wish to leave for future development, see below), this neglects the possibility that susceptibles can have contact with several infecteds in a smaller population or more structured contact network. That could be modelled by a force of infection, which would more realistically describe contraction of the generation interval during the peak of the outbreak, and provide estimates for relevant quantities such as reproduction ratios [6]. However, it requires information about uninfected susceptibles in the same population and a more complicated transmission model, which is a significant disadvantage when it comes to general applicability, one of our primary aims. More importantly, for transmission tree inference it does not seem to be a problem: the analyses of the published simulations were almost as accurate as in the original publication [19], and these simulations were in very small populations with almost all hosts infected.

The role of the within-host model is to integrate over all possible phylogenetic mini-trees and mutation events within the hosts. Therefore, the sometimes less efficient mixing of the within-host growth rate *r* (small ESS) is not problematic as it does not prevent good mixing of the tree topology. The role of the substitution model is to explain the genetic diversity in the data, through the likelihood of the genetic data (Eq (8)). We have used a wide prior on the mutation rate *μ*, and assumed a homogeneous site model. In principle these choices can easily be made more general in the same MCMC framework, but we have found that on the time scale of outbreaks, the likelyhood in very much dependent on the number of mutations in the tree (parsimony score). The method finds the phylogenetic tree with fewest mutations and matches the posterior *μ* to the number of mutations and the sum of the tree’s branch lengths, resulting in accurate estimates with both simulated and actual data. Allowing different rates for different sites (or for transversion vs. transition) will not change this, and will mainly result in more mutation rate estimates.

With two exceptions, the parsimony scores of posterior tree samples were always equal to the number of SNPs in the datasets (the minimum possible). The first exception is the set of *Fast Clock* published simulations, which had so many SNPs that many of the same mutations had occurred in parallel. The second exception is the H7N7 dataset. In that case, the analyses of the three genes separately resulted in parsimony scores with 6-12 (6%-13%) more mutations than the number of SNPs, whereas the analysis of all genes together resulted in a parsimony score of 313 (median) to explain only 257 SNPs, a surplus of 56 mutations (18%). The results for separate genes could indicate positive selection, confirming the analysis by Bataille et al [25], who even identified specific mutations that had occurred multiple times. The even higher discrepancy for the combined analysis is suggestive of reassortment events, also recognised by Bataille et al [25].

The proposed method and implementation opens perspectives for further extending the methodology to reconstruct phylogenetic and transmission trees from pathogen sequence data. One possible set of extensions arises from changes to the models embedded in our method, to include additional aspects of outbreak dynamics. For instance, the generation time distribution (infectiousness curve) could be made dependent on the sampling interval, which may be relevant for the MRSA outbreak analysis in which the two health-care workers may have transmitted the bacterium until late after infection. This dependence is implicit in methods in which transmission is modelled more mechanistically (e.g. [12, 13, 19]), but we chose not to do that to keep the model more generic. Another important extension would be to relax the assumption of a complete bottleneck at transmission; the bottleneck may be larger in reality [39, 40] and it has has previously been relaxed by looking at transmission pairs [41] or modelling it as separate transmission events [15], but not yet in a timed transmission tree. In our model, this would mean that a host can carry multiple phylogenetic mini-trees, rooted at the same infection time to the same infector. A third extenstion would be to include the possibility of reassortment of genes within a host, primarily motivated by the results of the H7N7 analysis. This may be done by modelling the coalescent process within hosts, the phylogenetic mini-trees, differently for different genes, but constrained by a single transmission tree. Finally, it would be possible to allow for multiple index cases, which may play a role in open populations with possible re-introductions (as in the MRSA setting), or when only a subset of a large epidemic is analysed (the FMD2001 dataset). This is implemented in models using genetic models based on pairwise genetic distances [7, 13], but is considered a major challenge with a coalescent model [42]. Multiple index cases could also reflect unobserved hosts in the outbreak itself, recently addressed by Didelot et al [43] in their two-step approach of first inferring a phylogenetic and then a transmission tree.

A second type of extension would stem from incorporating additional data. An example is the use of data that make particular transmissions more or less likely, such as contact tracing data, or censoring times for infection times per host or transmission times between sets of hosts, motivated by the MRSA dataset in which admission and discharge days are known for each patient. Sampling of infection times and infectors could be constrained by these additional data (as in [19, 27]). Another example is the use of spatial data in combination with a spatial transmission kernel, so that the likelihood of infectors includes a distance-dependency, a possible extension motivated by the FMD and H7N7 analyses (as in [24, 27]). A third example is the use of host characteristics to model infectivity as a function of covariates. With the MRSA data, it would then be possible to test for increased infectivity of the health-care workers, or to test for differences in transmissibility in the three wards. In general, the use of additional host data would make dealing with hosts for which a sequence is not available less problematic: the method currently can include these hosts, but without additional data their role is unclear and they are often placed at the end of transmission chains in consensus trees (Fig 2b, Fig 3).

## Methods

### Data

We developed our model for fully observed outbreaks of size *n* hosts. Data consist of the sampling times **S** and DNA sequences **G**, which means that for each host *i* we know the time of sampling or diagnosis *S_i_* and the sequence *G_i_* associated with the sampling time. It is not necessary to have a sequence for each host.

We illustrate the method with the following five datasets from earlier publications (all in S3 Data):

1. Tuberculosis *(Mycobacterium tuberculosis*, Mtb). This dataset was analysed by Didelot et al [11]. It consists of 33 Mtb cases in a population of drug users (approximate population size 400), with samples collected in a 2.5 years time frame. The 20 SNPs were part of a 4.4 Mbp long sequence. Analysis of this dataset tests the performance of this method in an outbreak with relatively few cases in a large population.
2. Methicillin-resistant *Staphylococcus aureus* (MRSA). This is the dataset from Nubel et al [21], with 36 MRSA cases in a neonatal ICU sampled within a time period of 7 months. Sampling dates were available for all cases, but sequences only for 28 cases, revealing 26 SNPs in the non-repetitive core genome of 2.7 Mbp. Analysis of this dataset tests for the performance of this method in an outbreak in a small population, including cases without sequence.
3. Foot-and-mouth disease (FMD2001). This is the dataset from Cottam et al [22] also analysed by several others [12, 24], with 15 infected premises within a time period of 2 months. Sequences were available for all cases, with 85 SNPs among 8196 nucleotides. Analysis of this dataset and the next tests for the performance of this method in a small completely sampled outbreak in a large population and allows comparison of the estimated transmission tree to earlier results.
4. Foot-and-mouth disease (FMD2007). This is the dataset from Cottam et al [23], also analysed by Morelli [24], with 11 infected premises within a time period of 2 months. Sequences were available for all cases, with 27 SNPs among 8176 nucleotides
5. H7N7 avian influenza (H7N7). This dataset has been analysed by several authors [19, 25-27], and consists of 241 poultry farms in a time period of about 2.5 months. Sequences of the HA, NA, and PB2 genes were available on GISAID for 228 farms, with associated sampling dates. The total number of SNPs was 257 in 5541 nucleotides. For the 13 unsampled farms we used the culling date minus 2 days as the observation day (the mean sampling-to-culling interval was 2.4 days in the 228 sampled farms). We analysed the data for the three genes separately, and combined. To inform a prior distribution for the interval from infection to sampling, we used estimated infection times from Boender et al [35]. Analysis of this dataset tests for the performance of this method in a large outbreak, including cases without sequence.

### The model and likelihood

The model describes the spread of an infectious pathogen in a population through contact transmission, the dynamics of the pathogen within the infected hosts, and mutation in the DNA or RNA of that pathogen. Furthermore, it describes how these dynamics are observed through sampling of pathogens in infected hosts. We infer the transmission tree and parameters describing the relevant processes by a Bayesian analysis, using Markov-Chain Monte Carlo (MCMC) to obtain samples from the posterior distributions of model parameters and transmission trees (infectors and infection times of all cases). We first introduce the models and likelihood functions; then we explain how we update the transmission trees and parameters in the MCMC chain.

The posterior distribution is given by

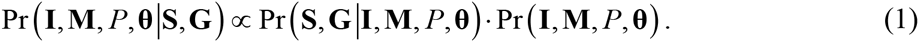

Equation (1) is the probability for the unobserved infection times **I**, infectors **M**, phylogenetic tree *P*, and model parameters **θ**, given the data (sampling times and sequences). The posterior probability can be split up into separate likelihood terms representing the four processes, times a prior probability for the parameters (see S2 Methods):

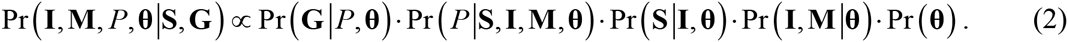

We now introduce the four models, the associated likelihoods, and prior distributions for associated parameters.

#### Transmission

We assume that the outbreak started with a single case. Each case produced secondary cases at random generation intervals after their own infection (Gamma distribution with shape *a_G_* and mean *m_G_*). We consider that all untimed transmission tree topologies are equally likely, so that the probability of the transmission tree only depends on its timing. The outbreak is described by the vectors I and M with infection times *I_i_* and infectors *M_i_* for all numbered cases *i*. The infector of the index case is 0. The likelihood is the product of probability densities (*d*_Γ(*a_G_,m_G_*)_ (·)) of all generation times in the outbreak:

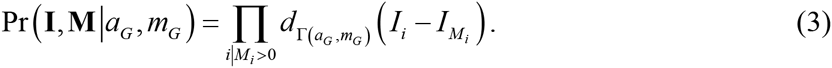

#### Sampling

We assume that all cases are observed and sampled at random times after they were infected, according to a Gamma distribution with shape *a_S_* and mean *m_S_*. Transmission and sampling are independent, so transmission can take place after sampling, and a case can be sampled earlier than its infector. The likelihood is the product of probability densities of all sampling intervals in the outbreak:

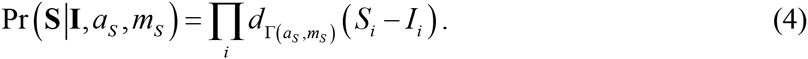

#### Within-host dynamics

The main function of the within-host model is to allow for a stochastic coalescent process within the host. Each host *i* harbours its own phylogenetic mini-tree *P_i_*, with the tips being the transmission and sampling events, and the root being the time of infection. Thus, the likelihood is the product of all likelihoods per host:

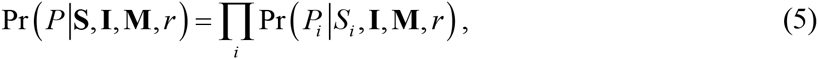

in which *r* is the parameter describing the within-host dynamics (see below). The dependency on all infection times and infectors remains for the mini-trees, because these determine the transmission times with host *i* as infector.

Going backwards in time, coalescence between any pair of lineages within a host takes place at rate 1/*w*(*τ, r*), where *w*(*τ, r*) = *rτ* denotes the linearly increasing within-host pathogen population size at (forward) time *τ* since infection of the host. With this particular function coalescent nodes tend to be close to the transmission events if *r* is small, whereas they tend to be soon after infection of the infector if *r* is large. This function also naturally results in only one lineage at the time of infection (complete transmission bottleneck), as the coalescence rate goes to infinity near the time of infection.

In the complete phylogenetic tree *P*, three types of nodes *x* are distinguished: nodes *x* = 1…*n* are the sampling nodes of the corresponding hosts *i* = 1…*n*, i.e. the tips of the tree at which sampling took place; nodes *x* = *n*+1…2*n*-1 are the coalescent nodes; nodes *x* = 2*n*…3*n*-1 are the transmission nodes, i.e. the points in the tree at which a lineage goes from one host to the next. By *h_x_* we identify the host in which node *x* resides; for transmission nodes it identifies the primary host (infector). The mini-tree *P_i_* is the set of nodes within host *i*, and *τ_x_* is the time of node *x* since infection of host *h_x_*. Let *L_i_* (*τ*) denote the number of lineages in host *i* at time *τ* since infection:

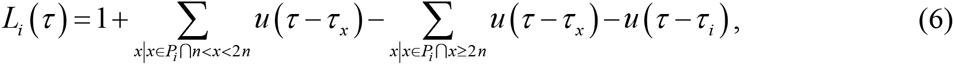

in which *u*(*τ*) is the heaviside step function, i.e. *u* (*τ*) = 0 if *τ* < 0, and *u* (*τ*) = 1 if *τ* ≥ 0. In other words, *L_t_* (0) = 1 by definition because of the complete transmission bottleneck, and then it increases by 1 at each coalescent node and decreases by 1 at each transmission event and at sampling. The likelihood for each mini-tree can then be written as

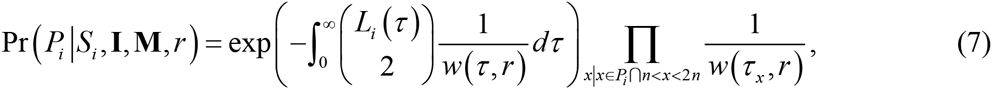

with 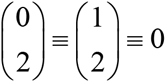. The first term is the probability to have no coalescent events during the intervals in which there are two or more lineages, the second term is the product of coalescent rates at the coalescent nodes.

#### Mutation

We use a single fixed mutation rate *μ* for all sites, with mutation resulting in any of the four nucleotides with equal probability (Jukes-Cantor). This parameterisation means that the effective rate of nucleotide change is 0.75*μ*. Given the phylogenetic tree, this results in the likelihood:

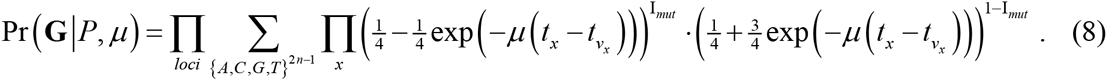

Here, we multiply over all coalescent and transmission nodes *x*, which occur at time *t_x_* and have parent node *v_x_*; I_*mut*_ indicates if a mutation occurred on the branch between *x* and *v_x_*. The likelihood is calculated using Felsenstein’s pruning algorithm [44].

#### Prior distributions

Here we describe our general choice of prior distributions, not the particular parameterization in our analyses (Section *Evaluating the method*). We chose fixed values for *a_G_* and *a_S_*, the shape parameters of generation and sampling intervals. For their means *m_G_* and *m_S_*, we used prior distributions with means *μ_G_* and *μ_S_* and standard deviations *σ_G_* and *σ_S_*, which are translated into Gamma-distributed priors for rate parameters *b_G_* = *a_G_*/*m_G_* and *b_S_* = *a_S_*/*m_S_*, distributed as Γ(*a*_0,*G*_, *b*_0,*G*_) and Γ(*a*_0,*S*_, *b*_0,*S*_) (see S2 Methods). For the slope *r* of the within-host growth model, we chose a Gamma-distributed prior with shape and rate *a_r_* and *b_r_*. We chose log(*μ*) to have a uniform (improper) prior distribution, equivalent to Pr(*μ*) ~ 1/*μ*.

### Inference method

We use Bayesian statistics to infer transmission trees and estimate the model parameters from the data, and MCMC methods to obtain samples from the posterior distribution. The procedure is implemented as a package in R (*phybreak*), which can be downloaded from GitHub (www.github.com/donkeyshot/phybreak). The package also contains functions to simulate data, and to summarize the MCMC output.

The main novelty of our method lies in the proposal steps for the phylogenetic and transmission trees, used to generate the MCMC chain. It makes use of the hierarchical tree perspective, in which the phylogenetic tree is described as a collection of phylogenetic mini-trees (one for each host), connected through the transmission tree. Most proposals are done by taking one host, changing its position in the transmission tree, and simulating the phylogenetic mini-trees in the hosts involved in that change. In a second type of proposal, the transmission tree is changed while keeping the phylogenetic tree intact.

Initialization of the MCMC chain requires initial values for the six model parameters (*a_G_, m_G_, a_S_, m_S_, r*, and *μ*). The transmission tree is initialized by generating an infection time for each host (sampling day minus random sampling interval). The first infected host is the index case, and for the remaining hosts an infector is randomly chosen, weighed by the density of the generation time distribution. Finally, the phylogenetic mini-trees in each host are simulated according to the coalescent model and combined with one another to create a complete phylogenetic tree.

Each MCMC iteration cycle starts with updates of the transmission and phylogenetic trees, followed by updates of the model parameters. To start with the latter, the parameters *m_S_* and *m_G_* are directly sampled from their posterior distribution given the current infection times and transmission tree (Gibbs update). This is done by sampling the rate parameters *b_S_* and *b_G_*, which were given conjugate prior distributions (see above). If 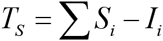 is the sum of *n* sampling intervals in the tree, *a*_0,*S*_ and *b*_0,*S*_ are the shape and rate of the prior distribution for *b_S_*, then a new posterior value is drawn as

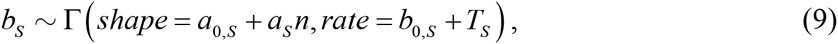

from which *m_S_* is calculated as *a_S_*/*b_S_*. Posterior values for *m_G_* are drawn from a similar distribution, with 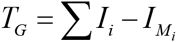. the sum of *n* – 1 generation intervals. The parameters *r* and *μ* are updated by Metropolis-Hastings sampling; proposals *r*’ and *μ*’ are generated from lognormal distributions *LN*(*r,σ_r_*) and *LN*(*μ,σ_μ_*), i.e. with current values as mean. The standard deviations are calculated based on the expected variance of the target distributions, given the outbreak size for *σ_r_*, and number of SNPs for *σ_μ_* (see S2 Methods).

#### Updating the phylogenetic and transmission trees

The phylogenetic and transmission trees, described by the unobserved variables **Z** = {**I, M**, *P*}, are updated by proposing a new tree with proposal density *H*(**Z’|Z, S, θ**), and accepting with Metropolis-Hastings probability (using Eq (1)) *α*,

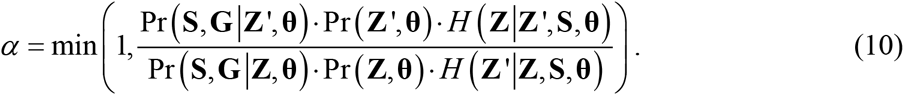

Per MCMC iteration cycle, *n* proposals are done with each host as a focal host once, in random order. Each proposal starts by taking a focal host *i*, drawing a sampling interval 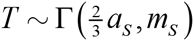 from a Gamma distribution with shape parameter 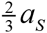 and mean *m_S_*, and calculating a preliminary proposal for the infection time *I_i_*’ = *S_i_* – *T*. Based on this preliminary proposal, the topology of the transmission tree is changed (see below), and in most cases the phylogenetic tree as well (80% probability). However, we also allowed for proposal steps without changing the phylogenetic tree (20% probability); this greatly improves mixing of the MCMC chain if there are many SNPs, which more or less fixes the phylogenetic tree topology. The 80%-20% distribution for the two types of proposal was not optimized but chosen such that mixing of the phylogenetic tree is only limitedly less efficient than without the second type of proposal (keeping the phylogenetic tree fixed).

#### Proposals for changes in transmission and phylogenetic trees

Here we describe how changes in the transmission and phylogenetic trees are proposed for six different situations, based on the preliminary proposal for the infection time *I_i_*’ and on whether the index case is involved. Fig 5 shows the proposed changes. More detail on the proposal distribution and calculation of acceptance probability is given in the S2 Methods.

A. The focal host *i* is index case, and the preliminary *I_i_*’ is before the first transmission event. In that case, the infection time of host *i* becomes *I_i_*’, and no topological changes are made in the transmission tree (Fig 5A).
B. The focal host *i* is index case, and the preliminary *I_i_*’ is after the first transmission event, but before host *i*’s second transmission event, if there is any. In that case, the infection time of host *i* becomes *I_i_*’, and host *i*’s first infectee becomes index case, transmitting to *i* (Fig 5B).
C. The focal host *i* is index case, and the preliminary *I_i_*’ is after host *i*’s second transmission event, if there is any. In that case, the infection times of host *i* and its first infectee are switched, and host *i*’s first infectee becomes index case. They may or may not exchange infectees, with 50% probability (Fig 5C).
D. The focal host *i* is not index case, and the preliminary *I_i_*’ is before infection of the index case. In that case, the infection time of host *i* becomes *I_i_*’, and host *i* becomes index case, transmitting to the original index case (Fig 5D).
E. The focal host *i* is not index case, and the preliminary *I_i_*’ is after infection of the index case, but before host *i*’s first transmission event. In that case, the infection time of host *i* becomes *I_i_*’, and a new infector is proposed from all hosts infected before *I_i_*’ (Fig 5E).
F. The focal host *i* is not index case, and the preliminary *I_i_*’ is after host i’s first transmission event. In that case, the infection times of host *i* and its first infectee are switched, as well as their position in the transmission tree. They may or may not exchange infectees, with 50% probability (Fig 5F).

**Fig 5.**
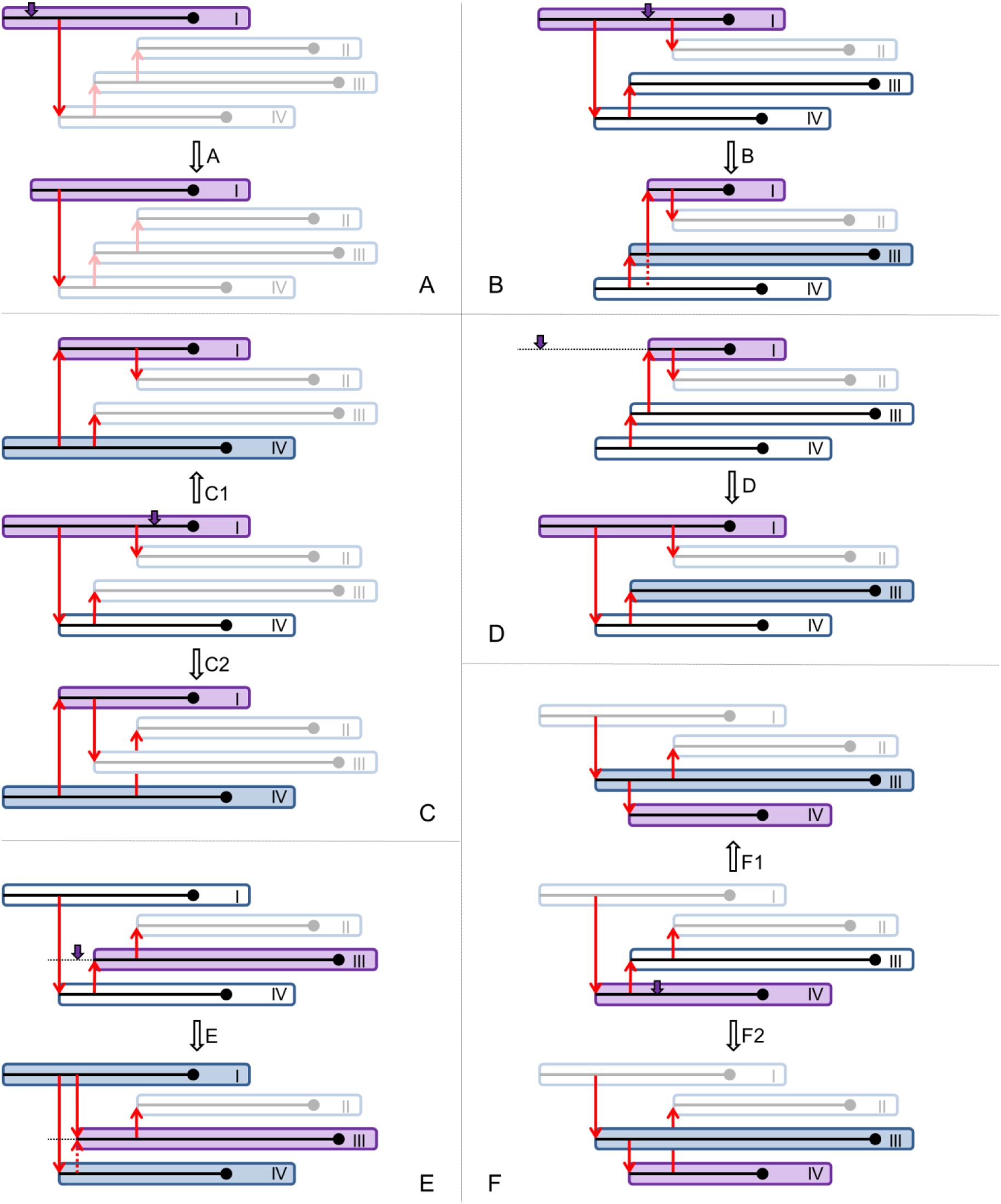
Graphics depicting proposal steps A-F for new transmission and phylogenetic trees. In panels A, B, D, and E, the initial situation is at the top, and the proposal below. In panels C and F, the initial situation is in the middle, and two alternative proposal above and below. Every panel shows an outbreak with four hosts, with red arrows indicating transmission: the purple host is the focal host, with the purple arrow indicating the proposal for the new infection time *I_i_*’; filled hosts have a new phylogenetic mini-tree proposed; greyed-out hosts do not play a role in the proposal. (A) the focal host is the index case, and *I_i_*’ is before the first transmission event; (B) the focal host is the index case, and *I_i_*’ is after the first, but before the second secondary case; (C) the focal host is the index case and *I_i_*’ is after his second secondary case; (D) the focal host is not the index case and *I_i_*’ is before infection of the index case; (E) the focal host is not the index case and *I_i_* is before his first secondary case; (F) the focal host is not the index case and *I_i_*’ is after his first secondary case.

Each change in the transmission tree is followed by proposing new phylogenetic mini-trees for all hosts involved, i.e. if their infection time was changed or transmission nodes were added or removed (grey hosts in Fig 5).

#### Proposals for changes in the transmission tree only

Here we describe how changes in the transmission tree are proposed without changing the phylogenetic tree, based on the preliminary *I_i_*’ and on whether the index case is involved. Fig 6 shows the proposed changes. More detail on the proposal distribution and calculation of acceptance probability is given in the S2 Methods.

G. The focal host *i* is the index case. If the preliminary *I_i_*’ is before the first coalescence node, the infection time of host *i* becomes *I_i_*’, and no changes are made in the transmission and phylogenetic trees. If the preliminary *I_i_*’ is after the first coalescence node, the proposal is rejected.
H. The focal host *i* is not the index case, and the preliminary *I_i_*’ is after the most recent common ancestor (MRCA) of the samples of host *i* and his infector *j*, which is a coalescent node in infector *j*. In that case, the infection time of host *i* becomes *I_i_*’, and infectees may move from host *i* to infector *j* or vice versa (Fig 6A).
I. The focal host *i* is not the index case, but his infector *j* is the index case, and the preliminary *I_i_* is before the MRCA of the samples of host *i* and his infector *j*. In that case, an infection time *I_j_*’ is proposed for the infector *j*. If *I_j_*’ is after the MRCA, the infection time of the infector *j* becomes *I_j_*’, and the infection time of host *i* becomes the original infection time of his infector *j*. Infectees may move from host *i* to infector *j* or vice versa (Fig 6B). If *I_j_*’ is before the MRCA, the proposal is rejected.
J. The focal host *i* is not the index case, and neither is his infector *j*, and the preliminary *I_i_*’ is before the MRCA of the samples of host *i* and his infector *j*, but after the MRCA of the samples of host *i* and infector *j*’s infector. In that case, an infection time *I_j_*’ is proposed for the infector *j*. If *I_j_*’ is after the MRCA, the infection time of the infector *j* becomes *I_i_*’, and the infection time of host *i* becomes *I_i_*’. Infectees may move between host *i*, infector *j*, and infector *j*’s infector (Fig 6C). If *I_j_*’ is before the MRCA of host *i* and infector *j*, or *I_i_*’ is before the MRCA of host *i* and infector *j*’s infector, the proposal is rejected.

**Fig 6.**
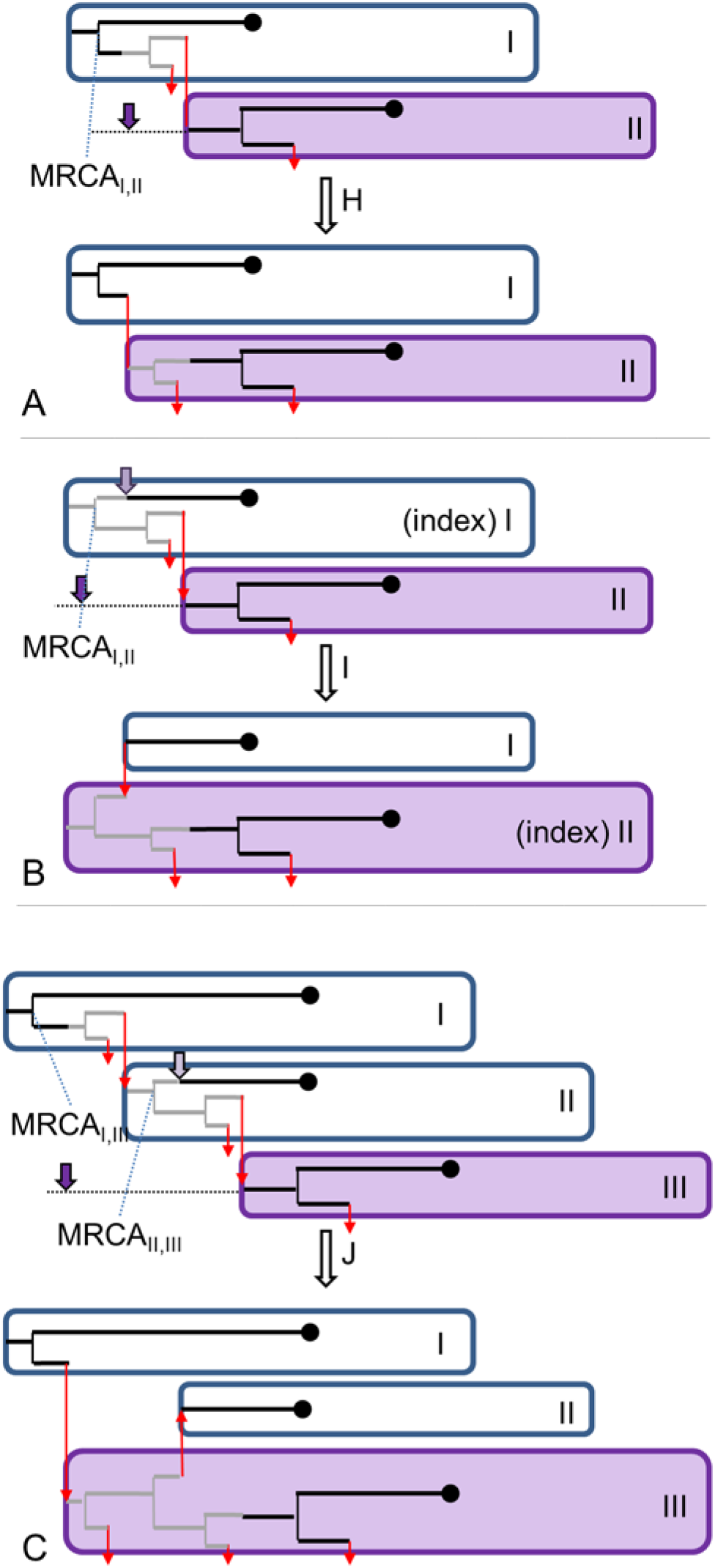
Graphics depicting proposal steps H-J for new transmission trees, keeping the phylogenetic tree unchanged. In all panels, the initial situation is at the top, and the proposal below. Every panel shows part of an outbreak, with red arrows indicating transmission to depicted or undepicted hosts. Only in panel B host I must be the index case. The purple host is the focal host, with the dark purple arrow indicating the proposal for the new infection time *I_i_*’; the light purple arrow in panels B and C indicate the proposal for the new infection time *I_j_*’ of the focal host’s infector. The grey parts of the phylogenetic tree are moved between the hosts. (A) the focal host is not the index case, and *I_i_*’ is after MRCA_I,II_ of the focal host and his infector; (B) the focal host is not the index case, and *I_i_*’ is before MRCA_I,II_ of the focal host and his infector (the index case), and *I_j_*’ is after MRCA_I,II_; (C) the focal host is not the index case, and *I_i_*’ is before MRCA_II,III_ of the focal host and his infector, but after the MRCA_I,III_ of the focal host and his infector’s infector; also, *I_j_*’ is after MRCA_II,III_.

### Evaluating the method

We took three approaches to evaluate our method: analysis of newly simulated data, analysis of published simulated data [19], and analysis of published observed data. When not specified, the following parameter settings and priors were used: shape parameters for sampling and generation interval distributions *a_S_* = *a_G_* = 3, uninformative priors for mean sampling and generation intervals with *μ_S_* = *μ_G_* = 1 and *σ_S_* = *σ_G_* = ∞, and an uninformative prior for within-host growth parameter *r* with *a_r_* = *b_r_* = 1. The prior for log(*μ*) (mutation rate) is always uniform.

Analyses were done by two MCMC chains, in each taking 25,000 samples (25,000 MCMC cycles). Burn-ins were different: 2000 MCMC cycles for the newly simulated data, 10,000 for the published simulated data [19], and 5000 for the observed data. With the H7N7 data, five MCMC chains were run, with a burn-in of 5000 samples, followed by 25,000 samples.

#### Analysis of newly simulated data

Four outbreak scenarios were simulated, each replicated 25 times: outbreak sizes of 20 and 50 cases, each with *a_G_* = *a_S_* = 3, resulting in overlapping generations and cases sampled earlier than their infector, or *a_G_* = *a_S_* = 10, resulting in more discrete generations and cases mostly sampled in order of infection. Further, the mean generation and sampling intervals were *m_G_* = *m_S_* = 1 year, the mutation rate *μ* = 10^-4^ per year in a DNA sequence with 10^4^ sites resulting in a genome-wide mutation rate of 1 per year and a number of SNPs in the same order of magnitude as the outbreak size. For the within-host model we used *r* = 1 per year.

The transmission trees were simulated assuming populations of size 35 or 86 individuals and *R*_0_ = 1.5, corresponding to expected final outbreak sizes of about 20 and 50 [45], respectively. Simulations started with one infected individual. All individuals were assumed to be equally infectious, resulting in a Poisson-distributed number of contacts at times since infection drawn from the generation time distribution; these contacts were made with randomly selected individuals and resulted in transmissions if that individual had not been infected before. Simulations were repeated until 25 outbreaks were obtained of the desired size.

Given the infection times, sampling times were drawn, and phylogenetic mini-trees were simulated for each host. These were combined into one phylogenetic tree on which random mutation events were placed according to a Poisson process with rate 1. Each mutation event was randomly assigned to one site, and generated one of the four nucleotides with equal probabilities (reducing the effective mutation rate by 25%). By giving the root an arbitrary sequence, the sampled sequences were obtained by following the paths from root to sample and changing the nucleotides at the mutation events.

The simulated data (sampling times and sequences) were analysed with four sets of parameter settings:

- Reference: *a_G_* = 3, all other parameters at simulation value (except for μ);
- Informative Correct: *a_S_* at simulation value, informative prior for *m_S_* with *μ_S_* = 1 and *σ_S_* = 0.1;
- Uninformative: *a_S_* at simulation value;
- Informative Wrong: *a_S_* at simulation value, informative prior for *m_S_* with *μ_S_* = 2 and *σ_S_* = 0.1.

#### Analysis of published simulated data

We used two sets of 25 simulated outbreaks, identified as *Fast clock* and *Slow clock* in the original paper [19], in which full details on the simulations can be found. Briefly summarizing some characteristics, 50 hosts were placed on a grid and a spatial transmission model was run, with exponential transmission kernel. Outbreaks with fewer than 45 cases were discarded. An SEIR (susceptible – exposed – infectious – removed) transmission model was used, with fixed latent period of 2 days and normally distributed infectious period (mean(sd) of 10(1) days). Sampling occurred at the time of removal. Phylogenetic mini-trees were simulated using a logistic within-host growth model *w*(*τ*) = 0.1(1 + *e*^6^)/(1 + *e*^6–1.5*τ*^), starting at *w*(0) = 0.1, then growing to *w* (4) = 20.2 and going to *w*(∞) = 40.4. Sequences were generated with a 14,000 base pair genome and a mutation rate of 10^-5^ per site per day (*Slow clock*) or 5·10^-4^ per site per day (*Fast clock*). The *Slow Clock* resulted in a mean number of mutations of 0.14 per day, or 0.98 per mean generation time of 7 days (latent period plus half infectious period), equivalent to the rate used in the new simulations; the *Fast Clock* was 50 times as fast.

The simulated data (sampling times and sequences, not locations and removal times) were analysed with three levels of prior knowledge on the sampling interval distribution:

- Naive: default settings;
- Uninformative: *a_S_* = 144 (coefficient of variation of 0.083, as in the simulation);
- Informative: *a_S_* = 144, an informative prior for *m_S_* (*μ_S_* = 12, *σ_S_* = 1).

#### Analysis of published datasets

The published Mtb, FMD2001, and FMD2007 datasets were analysed with default settings. The MRSA data contained information on times between hospital entry and first positive sample for 32 patients. Because of their mean and standard deviation of 20 days, we analysed these data with different prior information on the sampling interval only: *a_S_* = 1, *μ_S_* = 15, *σ_S_* = 5. For the H7N7 outbreak data, infection times of the flocks had been estimated [35], from which the mean and standard deviation of the sampling interval was calculated (7.0 and 2.2 days). We used this to inform the sampling intervals with: *a_S_* = 10, *μ_S_* = 7, *σ_S_* = 0.5. Because transmission after culling is not possible, we also used a weak informative prior for the mean generation interval: *μ_G_* = 5, *σ_G_* = 1.

#### Performance and outcome measures

The aim of the method is to reconstruct outbreaks in terms of infection times of all hosts and the transmission tree. This requires good mixing of the MCMC chain, especially of infection times and infectors, and a useful method to summarize all sampled transmission trees into a consensus tree.

To test for good mixing, we used effective sample sizes (ESS, calculated with the coda package in R) to evaluate mixing of the parameters and infection times. There are no strict thresholds, but in BEAST, an ESS < 100 is considered too low, whereas an ESS > 200 is considered sufficient [46]. Mixing of the tree topology (infector per host) was evaluated as follows. To test for 200 independent samples, the chains were thinned by 250, giving 100 samples per chain. Then two Fisher’s exact tests were done for each host, the first to compare the posterior frequency distributions of infectors across the chains (100 infectors per chain), the second to test for independency of subsequent samples, i.e. autocorrelation, within the chains (198 pairs of infectors). We used the proportion of successful tests (i.e. *P* > 0.05) as a measure of mixing, expecting 95% successful tests with good mixing.

Two methods were used to make consensus transmission tree topologies (who infected whom), both based on the frequencies of infectors for each host among the 50,000 posterior trees. The support of host *j* being the infector of host *i* is defined as the proportion of posterior trees in which host *i* infected host *j*. The first consensus tree is the maximum parent credibility (MPC) tree [19], which is the tree among all posterior trees that has the highest product of infector supports. The second consensus tree is the tree constructed using an adaptation of Edmond’s algorithm, which starts by taking the infector with highest support for each host, and resolves cycles if there are any [20]. Because the actual algorithm requires prior choice of an index case, we adapted it by repeating the algorithm for all supported index cases, and selecting the tree with highest sum of posterior supports (the measure used in the algorithm itself).

Posterior infection times were summarized either outside the context of a consensus tree, i.e. based on all MCMC samples, or for a particular consensus tree, i.e. for each host based only on those samples in which the infector was the consensus infector. The latter is to improve consistency between topology and infection times, although even then consistency is not guaranteed. For plotting transmission trees only, we used the Edmond’s consensus tree; for plotting transmission and phylogenetic trees together, we used the MPC consensus tree, which comes with a consistent phylogenetic tree because it is one of the sampled trees.

## Acknowledgements

We wish to thank Matthew Hall for sharing his simulated data [19], and authors of the publications [21-23, 25] for publicly sharing their outbreak data.

## Supporting Information

**S1 Results. Tables with additional results on simulated data.**

**S2 Methods. Extensive treatment of model and MCMC updating steps.**

**S3 Data. Sequence data and sampling times of analysed actual datasets.**

